# Theoretical minimum uncertainty of modulation enhanced spinning disk confocal microscopy

**DOI:** 10.1101/2023.08.25.554835

**Authors:** Dylan Kalisvaart, Shih-Te Hung, Carlas S. Smith

**Author notes:** Electronic mail.

## Abstract

Modulation enhanced single-molecule localization microscopy (meSMLM), where emitters are sparsely activated with patterned illumination, increases the localization precision over SMLM. Furthermore, meSMLM improves the resolution over structured illumination microscopy while reducing the required amount of illumination patterns. These factors motivate enabling meSMLM in existing systems which employ patterned illumination intensity. Here, we introduce SpinFlux: modulation enhanced localization for spinning disk confocal microscopy. SpinFlux uses a spinning disk with pinholes in its illumination and emission paths, to illuminate select regions in the sample during each measurement. The resulting intensity-modulated emission signal is analyzed to localize emitters with improved precision. We derive a statistical image formation model for SpinFlux and we quantify the theoretical minimum uncertainty, in terms of the Cramér-Rao lower bound, for various illumination pattern configurations. We find that SpinFlux requires multiple patterns to improve the localization precision over SMLM, with the maximum improvement being 1.17 when using a single pattern. When using two pinholes on opposing sides of the emitter position, the *x*-localization precision can locally be improved 2.62-fold over SMLM, whereas the *y*-precision is improved by maximally a factor 1.12. When using pinholes in a triangular configuration around the emitter position, the localization precision is balanced over the *x*and *y*-directions at approximately a twofold local improvement over SMLM, at the cost of suboptimal precision in each individual direction. When doughnut-shaped illumination patterns, created with a phase mask in the illumination and emission paths, are used for SpinFlux, the local precision improvement over SMLM is increased 3.5-fold in the *x*- and *y*-directions. While localization on ISM data ideally results in an average global improvement of 1.48 over SMLM, or 2.10 with Fourier reweighting, SpinFlux is the method of choice for local refinements of the localization precision.

Why it matters: One of the main objectives of singlemolecule localization microscopy (SMLM) is to improve the precision with which single molecules can be localized. This has been successfully achieved through modulation enhanced SMLM, which uses patterned illumination to increase the information content of signal photons. However, this technique relies on setups with increased technical complexity over SMLM. With SpinFlux, we locally enable a two-to 3.5-fold precision improvement over singlemolecule localization microscopy, which can be achieved with only minor modifications to existing spinning disk confocal microscopy setups (e.g. a phase mask in the illumination and emission paths). In addition, our modeling framework enables evaluation of a wide variety of spinning disk setups and therefore paves the way for optimal spinning disk design.

## I. INTRODUCTION

Single-molecule localization microscopy (SMLM) increases the precision with which single molecules can be localized beyond the diffraction limit^1–3^. Methods in SMLM require sparse activation of single emitters, after which emitters can be localized sequentially with reduced uncertainty. In recent years, various modulation enhanced SMLM (meSMLM) methods were introduced, which increase the localization precision over SMLM by sparsely activating emitters with intensity-modulated illumination patterns^4^. As a result, information is added to the data about the relative position of the emitter with respect to the illumination patterns. meSMLM methods include SIMFLUX^5^, SIMPLE^6^ and repetitive optical selective exposure (ROSE)^7^, which use sinusoidally-shaped intensity patterns, and MINFLUX^8^ and RASTMIN^9,10^, which use a doughnut-shaped illumination pattern. Patterned illumination can also be used to improve axial resolution, for example with modulated localization (ModLoc)^11,12^ and ROSE-Z^13^, which use illumination with both axial and lateral structure. Additional improvements to the localization precision can be attained through iterative meSMLM^14,15^, where patterns are iteratively moved through the sample using prior information from earlier measurements, to improve the localization precision locally around single-emitters.

Specifically for SIMFLUX^5^, it has been shown that meSMLM with sinusoidal patterns improves the resolution over both SMLM and structured illumination microscopy (SIM)^16^. SIM uses nine sinusoidal patterns in total aligned on three lateral axes, and subsequent reconstruction results in at most a twofold resolution improvement over the diffraction limit. SIMFLUX on the other hand only uses six patterns in total aligned on two lateral axes, and subsequent localization results in a 2.4-fold maximum improvement of the localization precision over SMLM. Therefore, the combination of structured illumination with sparse localization in meSMLM can result in a better resolution over existing reconstruction approaches, while using less illumination patterns in the process. These factors motivate the incorporation of meSMLM in existing systems, in which image reconstruction instead of localization is the current state-of-the-art. A promising candidate system is spinning disk confocal microscopy (SDCM)^17–21^ (see Figure 1a). SDCM introduces a spinning disk with pinholes in the illumination and emission paths. Rapidly pulsing the excitation laser causes stroboscopic illumination of the sample with moving illumination foci. If used for image scanning microscopy (ISM)^22^, the fluorescent emission signal is recorded on an image detector. Subsequent reconstruction of the recorded images results in an expected resolution improvement of a factor two over diffraction limited imaging^18,19^. Recently, SDCM was used for PAINT- and STORM-based localization microscopy, where SMLM localization algorithms were used to localize emitters in raw camera data^20,21^. It is shown that this improves the detection rate and signal-to-background ratio compared to widefield SMLM at the cost of a reduced signal photon count, resulting in a localization precision that is at best comparable to that of SMLM^20^. However, these methods do not take the information contained in the illumination pattern into account, as one would do in meSMLM. In this text, we therefore develop a statistical image formation model, suited for modulation enhanced localization in SDCM (see Figure 1b). Our method, called SpinFlux, uses patterned illumination generated by a spinning disk to excite the sample. The resulting intensity-modulated emission signal is then described by our image formation model. To evaluate the potential localization precision improvements of SpinFlux, we need to study the information contained in a single pattern exposure, the localization precision obtained by sequential illumination with multiple patterns and the optimal pattern configuration to maximally improve the precision. To accomplish this, we calculate the theoretical minimum uncertainty of SpinFlux in terms of the Cramér-Rao lower bound (CRLB)^23,24^. The CRLB is often used in (me)SMLM to quantify the theoretical minimum uncertainty of localizations. Using the SpinFlux image formation model, we calculate the CRLB for various illumination pattern configurations. Based on the CRLB, we compare SpinFlux with SMLM and an idealized comparable localization approach, where isolated emitters are localized in ISM reconstructions with a factor 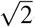reduction in the point spread function (PSF) width, or with a factor 2-reduction in case the ISM reconstructions are also Fourier reweighted (see Figure 1b). We find that SpinFlux requires at least two patterns to improve the localization precision over SMLM and ISM. We were able to locally minimize the localization uncertainty in one individual orientation, by using two pinholes with a radius of 3*σ*_PSF_ on opposing sides of the emitter position, where *σ*_PSF_ denotes the standard deviation of the emission point spread function. In this configuration, the *x*-localization precision can be improved 2.62-fold over SMLM when the patterns are spaced 4*σ*_PSF_ apart. For increasing separations until 4*σ*_PSF_, the information content of signal photons increases due to illumination with the low-intensity tails of the illumination patterns. However, when the separation increases above 4*σ*_PSF_, the loss of signal photons due to the windowing effect of the pinhole causes deterioration of the localization precision. While the two-pattern configuration maximizes the *x*precision, the *y*-precision is improved by maximally a factor 1.12. To circumvent this, we increase the amount of patterns used for SpinFlux. To maximize the localization precision, ideally pinholes are positioned in an equilateral triangle around the emitter position. This configuration balances the localization precision improvement over the *x*and *y*-directions, at the cost of suboptimal precision in each individual direction. In this case, the local precision improvement in each direction is approximately two. By including a phase mask in the illumination and emission paths, arbitrary diffraction-limited illumination patterns can be used for SpinFlux. When using doughnut-shaped intensity profiles, SpinFlux is able to locally attain a 3.5-fold precision improvement over SMLM in the *x*and *y*-directions.

**FIG. 1.**
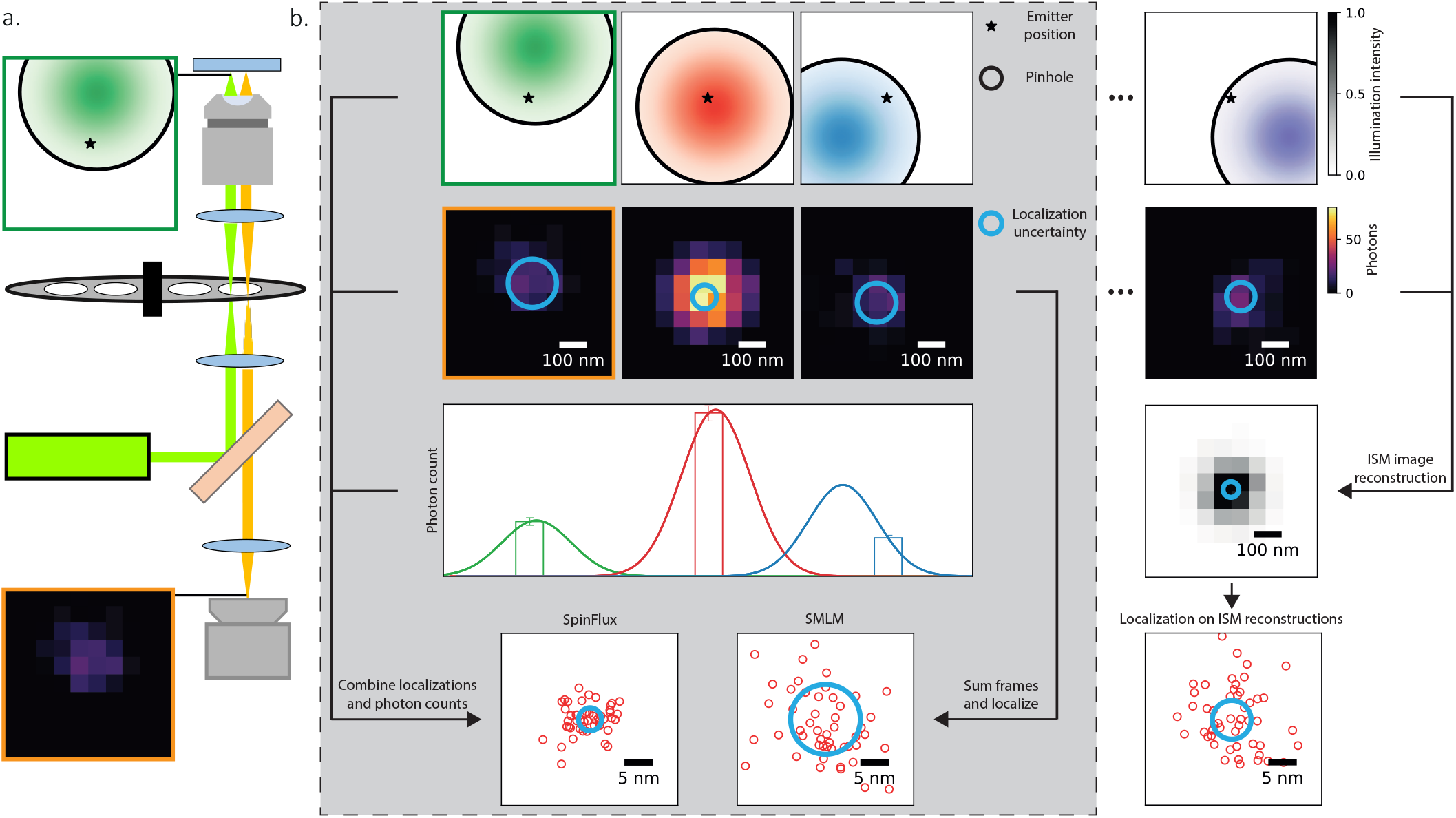
Schematic overview of SpinFlux image formation and analysis. (a) In SpinFlux, a spinning disk is placed in the illumination- and emission paths. This causes patterned illumination of emitters in the sample and subsequent windowing of the emission signal. Rapidly switching the laser on and off causes stroboscopic illumination of emitters in the sample with stationary illumination patterns. (b) SpinFlux obtains its localization precision improvement by merging localized emitter data with information about the relative distance between an illumination pattern and the emitter, derived from photon counts. In this way, it improves the localization precision over SMLM, which only uses localized emitter data and ignores pattern information. We compare SpinFlux with an idealized approach, in which first an ISM acquisition and reconstruction are performed, resulting in a 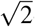-reduction of the PSF standard deviation, or a factor 2-reduction in case the ISM reconstructions are also Fourier reweighted. Afterwards, isolated emitters are localized in the ISM reconstruction

## II. METHODS

In SpinFlux localization (see Figure 1a), a spinning disk containing pinholes is placed in the illumination- and emission paths. The spinning disk is rotated, thereby sequentially moving illumination patterns over the sample. As in SDCM^19^, the excitation laser is rapidly switched on and off. Within the time frame where the laser is on, the spinning disk can be considered stationary. This causes stroboscopic illumination of emitters in the sample. Furthermore, the illumination has a non-uniform intensity profile over the field of view due to the spinning disk architecture. This causes patterned illumination of emitters in the sample, which in turn results in intensity modulation of the emission signal. Subsequently, the intensity-modulated emission signal is windowed by the same pinhole, after which the signal is imaged on a camera. The image analysis (see Figure 1b) consists of extracting localized emitters from the recordings, as well as retrieving the relative distance between the illumination pattern and emitter from the photon count. To evaluate the total amount of information that can be extracted from the measurements with this approach, we first develop an image formation model for SpinFlux. We subsequently use this model to calculate the theoretical minimum uncertainty of SpinFlux, in terms of the CRLB. The CRLB will allow us to quantify the maximum amount of information contained in each exposure with a single pattern. In turn, we use this to derive the localization precision that can be attained through sequential exposures with multiple patterns. In addition, we can explore how the pattern configuration, the pinhole radius and the mutual spacing between patterns affect the maximum localization precision.

### A. Model for SpinFlux image formation

To calculate the theoretical minimum uncertainty which can be attained with SpinFlux localization, we need a model to describe the amount of photons collected by a camera pixel. Existing models for (modulation-enhanced) single molecule localization microscopy^5,8,14,25,26^ do not suffice for this, as they do not include a pinhole in the illumination and emission paths. In this subsection, we therefore develop a statistical image formation model for SpinFlux. A detailed derivation of this model can be found in Supplementary Note 2.

For the image formation, we assume that pinholes are separated far enough on the spinning disk, such that only one pinhole can appear in a region of interest during each camera frame. This assumption is valid for the magnifications, pinhole sizes and pinhole separations in existing SDCM setups^19–21^. In line with this, we can assume that there is no crosstalk of emission signals between different pinholes. This allows us to describe the regions of interest on the camera frames as separate regions of interest from individual patterns.

We model the pinhole in the emission path as a circular window. In the absence of read-out noise, the measurements on each camera pixel can be described as independent realizations of a Poisson process^25^. For each pixel *i* with center coordinates (*x*_*i*_, *y*_*i*_) and for camera frame corresponding to illumination pattern *k*, the expected photon count *µ*_*i,k*_ after illumination through the pinhole with position (*x*_p,*k*_, *y*_p,*k*_) is described by (see Figure 1b, Supplementary Note 2):

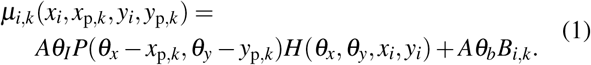

Here, (*θ*_*x*_, *θ*_*y*_) is the emitter position, *θ*_*I*_ is the expected signal photon count under maximum illumination and *θ*_*b*_ is the expected background photon count.

We model each illumination pattern *P*(*θ*_*x*_*−x*_p,*k*_, *θ*_*y*_ *−y*_p,*k*_) as a Gaussian PSF in the center of the pinhole, with standard deviation *σ*_illum_. Alternate illumination patterns can be generated by placing a phase mask in the illumination path. We therefore also include a model of the doughnut-shaped pattern from e.g. MINFLUX^8^, with a zero-intensity minimum at the center of the pinhole and standard deviation *σ*_illum_.

We model the emission PSF as a Gaussian, with standard deviation *σ*_PSF_. The term *H*(*θ*_*x*_, *θ*_*y*_, *x*_*i*_, *y*_*i*_) describes the discretized emission PSF after windowing by the pinhole (see Supplementary Note 2: Illumination and emission point spread functions).

In existing work on meSMLM, such as in MINFLUX^8^, it is assumed that meSMLM is able to record the same amount of signal photons as SMLM. This assumption allows benchmarking between methods on the same signal photon count. However, the assumption is not trivial, as additional illumination power or time is needed to exhaust the signal photon budget with non-maximum illumination intensity. Properly adjusting the illumination power to compensate for the reduced photon flux requires accurate prior knowledge about the emitter position, which is generally unavailable. Increasing the illumination time increases the probability of sample degradation. As such, we should include the possibility that meSMLM will not exhaust the signal photon budget in the image formation model.

The normalizing constant *A* describes how the signal photon budget is affected by non-maximum illumination intensity. This constant plays a vital role in benchmarking meSMLM (when the summed intensity over all patterns does not result in a uniform profile), as it gives a physical explanation of the fair signal photon count against which meSMLM should be compared^14^. Specifically when comparing meSMLM to SMLM, the normalization constant models whether meSMLM would have had recorded the same amount of signal photons as SMLM, despite the additional illumination power or time needed to do so. Results on the improvement of meSMLM compared to SMLM should thus only be given in the context of the normalizing constant *A*.

We choose *A* to model two scenarios (see Figure 1d, Supplementary Note 2: Multiple emission patterns). In the first scenario, we assume that the entire signal photon budget is exhausted after illumination with all patterns (aside from signal photons that are blocked by the spinning disk), disregarding the illumination power and time needed to accomplish this for each pattern. Here, *A* is inversely proportional to the summed illumination patterns. This scenario is consistent with the assumption used in e.g. MINFLUX^8^, stating that meSMLM will record the same amount of photons as SMLM. In the second scenario, the illumination power and time are constant for each pattern such that the total illumination power and time equal that of SMLM, even though this does not exhaust the signal photon budget for non-maximum illumination. Here, *A* is inversely proportional to the amount of illumination patterns *K*.

The constant *B*_*i,k*_ describes how the background is affected by illumination pattern *k*. As such, the term *Aθ*_*b*_*B*_*i,k*_ represents the effective background under patterned illumination. It depends on the camera pixel area, the pinhole area, the PSF and the illumination pattern, but not on the emitter position (see Supplementary Note 2: Effective background *B*_*i*_). In the analysis of e.g. MINFLUX^8^, the pattern-dependency of the background is neglected. We can incorporate this in our image formation model for SpinFlux, by modeling *B*_*i,k*_ as the intersection area between the camera pixel *i* and the approximation of pinhole *k* (see Supplementary Note 2: Pattern-independent background).

### B. Cramér-Rao lower bound

To quantify the theoretical minimum uncertainty of localizations, the Cramér-Rao lower bound is often used^23,24^. Under regularity conditions on the likelihood of the data^23^, the CRLB states that the estimator covariance 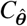 of any unbiased estimator 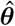 of the parameters ***θ*** satisfies the property that 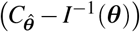 is positive semi-definite. Here, *I*(***θ***) is the Fisher information, of which entry (*u, v*) is described by:

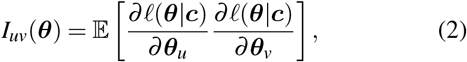

where *l* (***θ***|***c***) is the log-likelihood function given the recorded photon counts ***c*** on the camera pixels. The matrix *I*^*−*1^(***θ***) is the CRLB. Consequently, the diagonal of the CRLB bounds the estimator variance from below.

Specifically for SMLM, the CRLB is attained by the covariance of the maximum likelihood estimator for 100 or more signal photons^25^. As the localization uncertainty of the maximum likelihood estimator converges asymptotically to the CRLB^27,28^, we can also use the CRLB to investigate the theoretical minimum uncertainty of SpinFlux.

Using the image formation model from Equation (1), we can derive the Cramér-Rao lower bound for SpinFlux. When using *K* pinholes and a camera consisting of an array with *N*_pixels_ pixels, any entry (*u, v*) of the Fisher information is given by (see Supplementary Note 3):

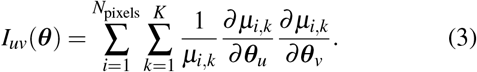

To evaluate Equation (3), the partial derivatives of the image formation model of Equation (1) with respect to the unknown parameters *θ*_*x*_, *θ*_*y*_, *θ*_*I*_ and *θ*_*b*_ need to be computed. Expressions for these partial derivatives are found in Supplementary Note 4.

### C. Simulations and parameter values

We sampled measurements from the image formation model and evaluated the CRLB using representative *in silico* experiments. The model parameters (see Supplementary Table 1) are considered to be representative of an SDCM experiment^20^.

To maximize the information contained in the Gaussian illumination and emission PSFs, we choose their standard deviations to be diffraction limited^29^. Specifically, we approximate the standard deviation of the illumination 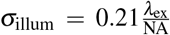 and the standard deviation of the PSF 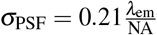. Here, *λ*_ex_ and *λ*_em_ respectively describe the excitation and emission wavelengths and NA is the numerical aperture.

Emitters are located in the center of the region of interest, consisting of 10 by 10 pixels. The pinhole was discretized on a mesh with *N*_M,*x*_, *N*_M,*y*_ = 100 pixels in each direction. For *N*_M,*x*_, *N*_M,*y*_ = 100 mesh pixels, the relative error in the CRLB caused by the discretized pinhole approximation is at most 0.02% (see Supplementary Figure 2).

## III. RESULTS AND DISCUSSION

The spinning disk of an SDCM setup can be designed with various pinhole sizes, spacing and arrangements^20^. Additionally, the rotation of the spinning disk gives additional freedom, as patterns and pinholes can appear arbitrarily close to each other via sequential illumination with a rotating spinning disk. For SpinFlux, this means that a wide variety of illumination pattern configurations can be created via the appropriate spinning disk and rotation speed.

Here, we compute the theoretical minimum localization uncertainty for three standard configurations. These pattern configurations can be realized either via simultaneous illumination with an appropriate spinning disk architecture or via sequential illumination with a rotating spinning disk. In Subsection III B, we do so for a single pattern and pinhole, akin to confocal microscopy. In Subsection III B, we establish localization on ISM reconstruction data as a benchmark for SpinFlux. In Subsection III C, we compute the CRLB for a two-pattern configuration where pinholes are separated by a distance *s* along the *x*-axis, resembling raster-like configurations of earlier work on meSMLM^9,10,14^. In Subsection III D, patterns and pinholes are arranged in an equilateral triangle configuration, similar to the configuration found in MINFLUX^8,15^. Variations of these configurations, such as those where configurations are rotated or where additional pinholes are added, can be found in Supplementary Figures 2 to 29. Supplementary Figures 31 to 42 show the effect of doughnut-shaped illumination patterns, generated via a phase mask in the illumination path (see Supplementary Figure 30), on the localization precision of SpinFlux.

As identified in Subsection II A, the normalizing constant *A* of Equation (1) needs to be chosen to describe how the signal photon budget is distributed over the individual patterns and to determine a fair photon count at which SpinFlux can be benchmarked against SMLM. To thoroughly discuss the improvement of SpinFlux, we evaluate the localization precision in three scenarios. In Figures 3 to 5 and Supplementary Figures 3 to 9 and 31 to 34, we calculate the theoretical minimum uncertainty for the scenario where the entire signal photon budget is exhausted after illumination with all patterns. Supplementary Figures 10 to 19 and 35 to 38 show the theoretical minimum uncertainty in case the illumination power and time are constant for each pattern. Lastly, Supplementary Figures 20 to 29 and 39 to 42 show the CRLB where the pattern-dependency of the background is neglected and where the entire signal photon budget is exhausted after illumination with all patterns.

### A. Localization on ISM reconstruction data

To benchmark the potential localization precision improvement of SpinFlux, we consider SMLM. Additionally, we consider an idealized comparable localization approach, consisting of localizing isolated emitters in ISM reconstruction data. In this approach, an ISM image is first acquired and reconstructed, resu<u>lt</u>ing in a reduction of the PSF width by at most a factor 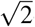. If the ISM image is subsequently Fourier reweighted^18^, the PSF width is reduced further by a total factor 2. Subsequently, individual emitters are localized in the ISM reconstruction data. For a signal photon count of 2000 photons per emitter and a background photon count of 8 photons per pixel, this ideally results in an improvement of 1.48 over SMLM without Fourier reweighting, or an improvement of 2.10 with Fourier reweighting (see Figure 2, Supplementary Note 1, Supplementary Figure 1). Note that as opposed to SpinFlux, this approach requires illumination with enough patterns (usually hundreds of patterns) to uniformly illuminate the sample.

**FIG. 2.**
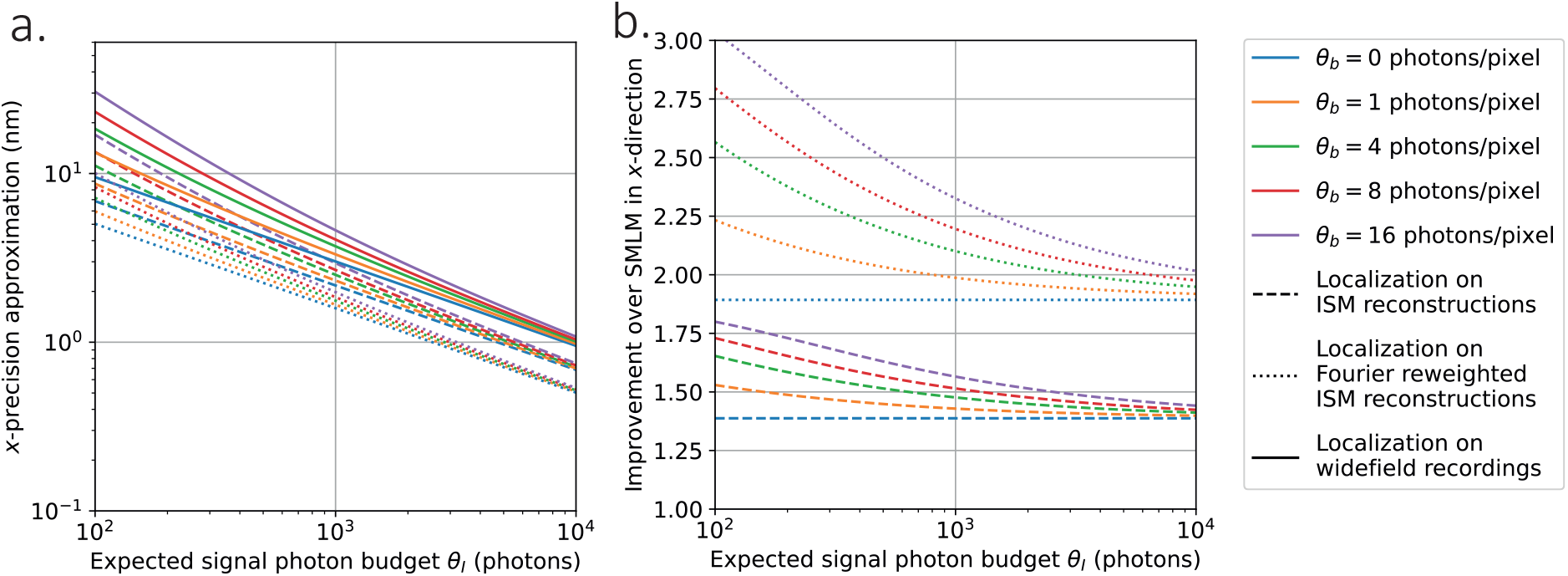
Approximation of the theoretical minimum localization uncertainty of single-molecule localization microscopy (SMLM) on data acquired from (Fourier reweighted) image scanning microscopy (ISM). A point spread function (PSF) standard deviation of 93.3 nm and a camera pixel size of 65 nm were used. **(a)** Approximate Cramér-Rao lower bound (CRLB) in *x*-direction as a function of the expected signal photon budget for varying values of the expected background photon count. **(b)** Improvement of the approximate CRLB over SMLM as a function of the expected signal photon budget for varying values of the expected background photon count.

### B. Single pattern configuration

In Figure 3, we evaluate the theoretical minimum uncertainty in case a single pinhole is used for illumination and emission, as illustrated in Figures 3a to 3c. Results are shown for the scenario where the entire signal photon budget is exhausted after illumination with all patterns.

**FIG. 3.**
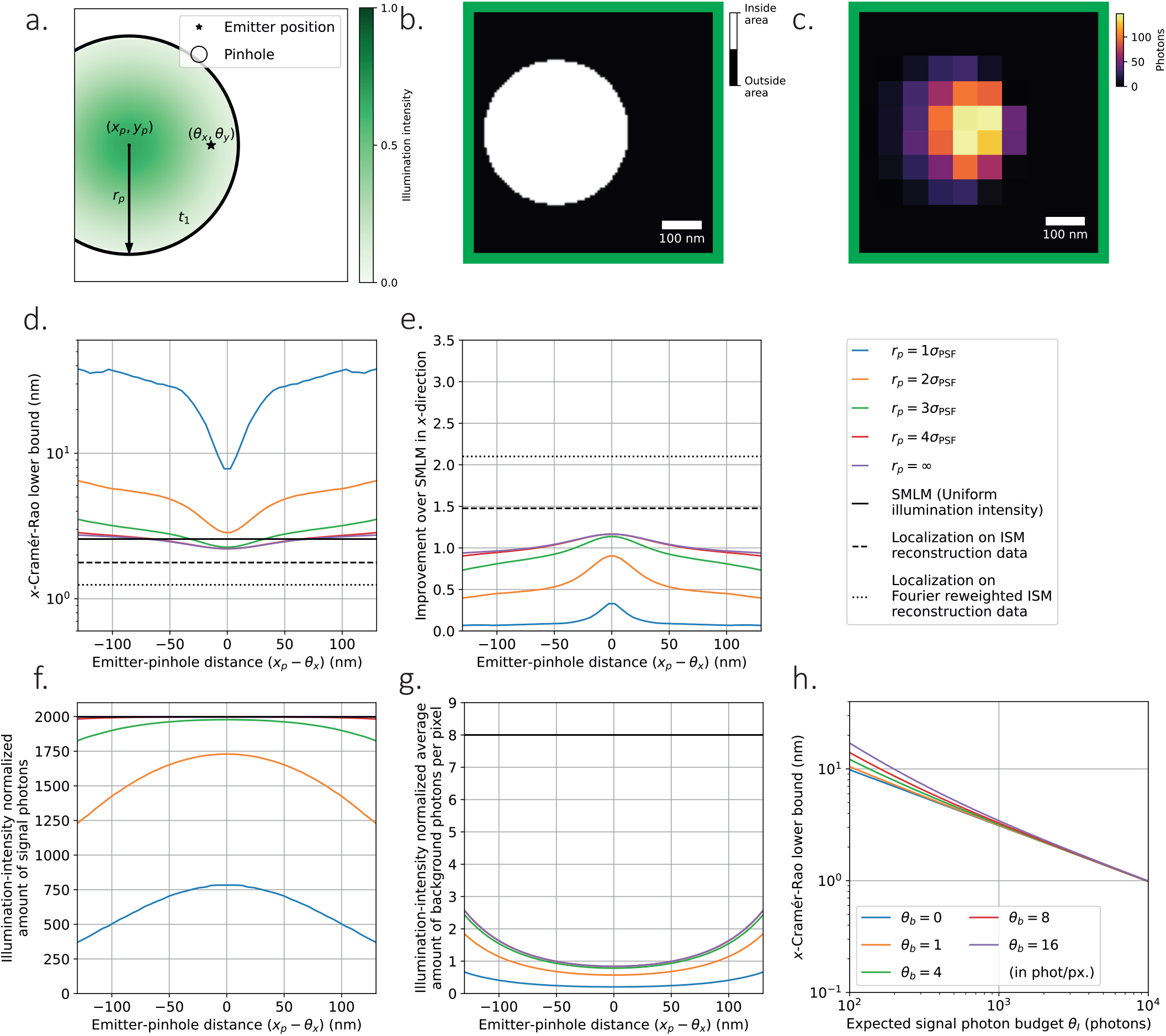
Theoretical minimum localization uncertainty of SpinFlux localization with one *x*-offset pinhole and pattern. In (c-g), 2000 expected signal photons and 8 expected background photons per pixel were used. Results are evaluated for the scenario where the entire signal photon budget is exhausted after illumination with the pattern (disregarding signal photons blocked by the spinning disk). **(a)** Schematic overview of SpinFlux localization with one pinhole with radius *r*_p_, centered at coordinates (*x*_p_, *y*_p_). In (d-g), the *x*-distance (*x*_p −_*θ*_*x*_) between the pinhole and the emitter is varied, where *y*_p_ = *θ*_*y*_. **(b)** Example of pinhole in the region of interest (650 ×650 nm). The pinhole radius *r*_p_ = 2*σ*_PSF_ was used. The pinhole mask was discretized with *N*_M,*x*_, *N*_M,*y*_ = 100 mesh pixels in each direction. **(c)** Example of fluorescent response in the region of interest, resulting from illumination and emission through the pinhole in (b). **(d)** Cramér-Rao lower bound (CRLB) in *x*-direction as a function of the emitter-pinhole *x*-distance. Simulations show SpinFlux with varying pinhole sizes and widefield single-molecule localization microscopy (SMLM). **(e)** Improvement of the SpinFlux CRLB over SMLM as a function of the emitter-pinhole *x*-distance for varying pinhole sizes. **(f)** Average amount of signal photons after compensation for non-maximum illumination intensity as a function of the emitter-pinhole *x*-distance, for SpinFlux with varying pinhole sizes and widefield single molecule localization microscopy (SMLM). **(g)** Average amount of background photons per pixel after compensation for non-maximum illumination intensity as a function of the emitter-pinhole *x*-distance, for SpinFlux with varying pinhole sizes and widefield single molecule localization microscopy (SMLM). **(h)** CRLB in *x*-direction as a function of the expected signal photon count for varying values of the expected background photon count. The pinhole radius *r*_p_ = 3*σ*_PSF_ was used and (*x*_p_, *y*_p_) = (*θ*_*x*_, *θ*_*y*_).

From Figures 3d and 3e, we see that the localization precision is optimal when the pinhole and pattern are centered directly on the emitter position. Without a pinhole, this results in an improvement of at most 1.17 over SMLM. For a pinhole with radius *r*_p_ = 4*σ*_PSF_, the difference with SMLM is negligible, indicating that the confocal effect of the pinhole has been lost. The improvement can thus be attributed to the effect of pattern-dependent background, as the background is reduced on camera pixels that are not located on the maximum of the Gaussian illumination pattern. This background reduction is visualized in Figure 3g, showing a 10.2-fold reduction in the average background count per pixel compared to SMLM for *r*_p_ = 4*σ*_PSF_ and *x*_p_ = *θ*_*x*_. A similar effect is shown in Supplementary Figure 10 in the scenario where the illumination power and time are fixed. The effect is not present in Supplementary Figure 20, which shows the results when the pattern-dependent background is neglected. In this case, no improvement over SMLM can be attained with a single pattern. Furthermore, the same effects are also present when considering different pattern *y*-coordinates, as shown in Supplementary Figures 3, 11 and 21 for each respective scenario.

For pinholes of radius *r*_p_ = 3*σ*_PSF_ and below, the localization precision deteriorates with respect to the no-pinhole case. Already for *r*_p_ = 2*σ*_PSF_, no position of the pinhole results in an improvement over SMLM. In these cases, the pinhole not only blocks background photons, but also signal photons carrying information about the emitter position. Figures 3f and 3g show that in the best case (for *x*_p_ = *θ*_*x*_) 248 signal photons are lost when going from *r*_p_ = 3*σ*_PSF_ to *r*_p_ = 2*σ*_PSF_, whereas the average background is reduced with only 0.21 photons per pixel. As such, more information about the emitter position is lost due to the loss of signal photons than that we gain by blocking background, resulting in a reduction of the improvement factor from 1.14 to 0.90. Similarly, moving the pinhole away from the emitter position blocks signal photons, thereby reducing the localization precision. For *r*_p_ = 3*σ*_PSF_, the improvement over SMLM goes from 1.14 at *x*_p_ = *θ*_*x*_ to 0.73 at a 130 nm distance between *x*_p_ and *θ*_*x*_. We find similar effects in the other scenarios under our consideration, as shown in Supplementary Figures 10 and 20. From this, we can conclude that bigger pinholes are in principle better for SpinFlux, as more information about the underlying signal is revealed through the bigger pinhole.

### C. Two-pattern configuration

In Figure 4, we evaluate the theoretical minimum uncertainty in case two pinholes are used for illumination and emission, which are separated in *x*-direction around focus coordinates (*x*_f_, *y*_f_) as illustrated in Figures 4a to 4c. Results are shown for the scenario where the entire signal photon budget is exhausted after illumination with all patterns. For these simulations, the pinhole radius was set to *r*_p_ = 3*σ*_PSF_ for both pinholes.

**FIG. 4.**
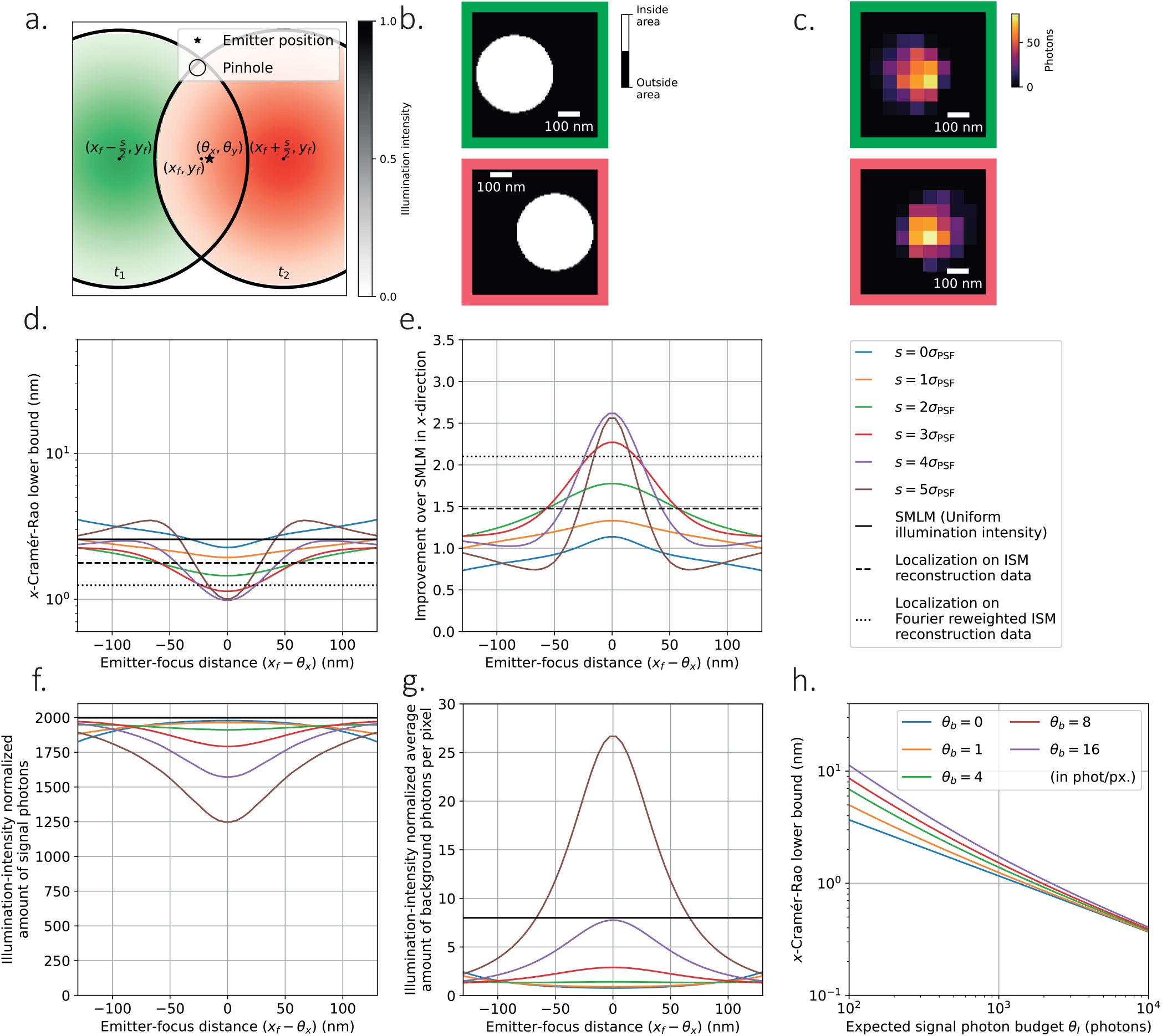
Theoretical minimum localization uncertainty of SpinFlux localization with two pinholes and patterns separated in the *x*-direction. In (c-g), 2000 expected signal photons and 8 expected background photons per pixel were used, with pinhole radius *r*_p_ = 3*σ*_PSF_. Results are evaluated for the scenario where the entire signal photon budget is exhausted after illumination with all patterns (disregarding signal photons blocked by the spinning disk). **(a)** Schematic overview of SpinFlux localization with two pinholes, separated in *x* and centered around the focus coordinates (*x*_f_, *y*_f_). In (d-g), the *x*-distance (*x*_f −_*θ*_*x*_) between the pattern focus and the emitter is varied, where *y*_f_ = *θ*_*y*_. **(b)** Example of pinholes in the region of interest (650 ×650 nm). The pinhole radius *r*_p_ = 2*σ*_PSF_ and pinhole separation *s* = 2*σ*_PSF_ were used. The pinhole masks were discretized with *N*_M,*x*_, *N*_M,*y*_ = 100 mesh pixels in each direction. **(c)** Example of fluorescent response in the region of interest, resulting from illumination and emission through each pinhole in (b). **(d)** Cramér-Rao lower bound (CRLB) in *x*-direction as a function of the emitter-focus *x*-distance. Simulations show SpinFlux with varying pinhole separations and widefield single molecule localization microscopy (SMLM). **(e)** Improvement of the SpinFlux CRLB over SMLM as a function of the emitter-focus *x*-distance for varying pinhole separations. **(f)** Average amount of signal photons after compensation for non-maximum illumination intensity as a function of the emitter-focus *x*-distance, for SpinFlux with varying pinhole separations and widefield single molecule localization microscopy (SMLM). **(g)** Average amount of background photons per pixel after compensation for non-maximum illumination intensity as a function of the emitter-focus *x*-distance, for SpinFlux with varying pinhole separations and widefield single molecule localization microscopy (SMLM). **(h)** CRLB in *x*-direction as a function of expected signal photon count for varying values of the expected background photon count. The pinhole radius *r*_p_ = 3*σ*_PSF_ and pinhole separation *s* = 4*σ*_PSF_ were used and (*x*_f_, *y*_f_) = (*θ*_*x*_, *θ*_*y*_).

From Figures 4d and 4e, we see that using multiple patterns is beneficial for SpinFlux, maximally resulting in a 2.62fold precision improvement over SMLM in the *x*-direction when using a pinhole separation *s* = 4*σ*_PSF_. This improvement descreases only moderately to 2.17 when the pattern *y*coordinate is moved 130 nm out of focus (see Supplementary Figure 4). When the illumination time and power are adjusted to exhaust the entire signal photon budget, the low-intensity tails of the Gaussian intensity profile increase the information content of signal photons, as these contain increased information about the relative position of the emitter with respect to the illumination pattern.

However, increasing the pinhole separation also reduces the region where SpinFlux improves over SMLM. For a pinhole separation *s* = 3*σ*_PSF_, the domain where SpinFlux improves over SMLM by at least a factor 1.2 spans 175 nm, whereas this domain spans 111 nm for *s* = 4*σ*_PSF_. In case the pinholes are not centered around the emitter position, one of the patterns takes more of the signal photon budget than the other. As such, highly informative signal photons carrying information from the tails of the Gaussian illumination pattern are traded in for lowly-informative photons coming from the center of the pattern. This is shown in Figure 4f: for a pinhole separation *s* = 4*σ*_PSF_, 1573 signal photons are collected in total when the *x*_f_ = *θ*_*x*_, with the remaining 427 photons being blocked by the spinning disk. When considering a 130 nm distance between *x*_f_ and *θ*_*x*_, 1956 signal photons are being collected in total as one pinhole has moved close to the emitter position. Yet these photons are lowly informative, resulting in a precision improvement of 1.09 over SMLM. For increasing separations, the relative difference in illumination intensity between non-centered patterns increases, thereby reducing the domain of improvement.

Furthermore, Figures 4d and 4e show that there is an optimal pinhole separation of *s* = 4*σ*_PSF_ for SpinFlux. When increasing the pinhole separation beyond this, the localization precision decreases again. This is caused by a combination of two factors. First of all, as shown in Figure 4f, the spinning disk blocks an increasing amount of signal photons for increasing pinhole separations, as the overlap between the pinhole and emission PSF is reduced. Between *s* = 4*σ*_PSF_ and *s* = 5*σ*_PSF_, the amount of signal photons is reduced by 324 when *x*_f_ = *θ*_*x*_. This effect is eliminated when the pinhole is removed, as shown in Supplementary Figure 5.

Secondly, increasing the pinhole separation results in illumination with the low-intensity tails of the Gaussian illumination patterns. As we exhaust the signal photon budget in this scenario and as the background is pattern-dependent, this results in an amplification of the background. Figure 4 shows that the average background count increases from 7.75 photons per pixel at *s* = 4*σ*_PSF_ to 26.7 photons per pixel at *s* = 5*σ*_PSF_. This effect is eliminated in the scenario where the illumination power and time are fixed, as shown in Supplementary Figures 12 to 15. Here, the background is not amplified as the signal photon budget does not need to be exhausted. However, due to the previously discussed effect of the signal photon count, the optimal separation is *s* = 2*σ*_PSF_ with an improvement of 1.31 over SMLM. Supplementary Figures 22 to 25 disregard the effect of pattern-dependent background entirely. Here, the pinhole separation *s* = 5*σ*_PSF_ achieves better localization precision than *s* = 4*σ*_PSF_, resulting in an improvement of 3.23 over SMLM.

Up until now, we have only considered the localization precision in the *x*-direction. To investigate how the two-pattern configuration of Figure 4 affects the *y*-precision, we equivalently consider the *x*-precision that can be obtained with the rotated pattern (see Supplementary Figures 6, 15, 25). From Supplementary Figure 6, we see that the *x*-precision for the rotated pattern results in negligible improvements or even reductions over SMLM if the entire signal photon budget is exhausted. Specifically for *s* = 4*σ*_PSF_, the improvement factor over SMLM is 0.83 when the patterns are perfectly centered around the emitter position, whereas the improvement increases to 1.12 when the distance between *y*_f_ and *θ*_*y*_ is 130 nm. Similar effects are found for the other scenarios under our consideration (see Supplementary Figures 15 and 25). From the equivalence, we can thus conclude that the two-pattern configuration of Figure 4 results in optimal *x*-precision, but the associated *y*-precision is diminished.

### D. Triangular pattern configuration

In Figure 5, we evaluate the theoretical minimum uncertainty in case multiple pinholes are used for illumination and emission in an equilateral triangle configuration, centered around focus coordinates (*x*_f_, *y*_f_) as illustrated in Figures 5a to 5c. Results are shown for the scenario where the entire signal photon budget is exhausted after illumination with all patterns. For these simulations, the pinhole radius was set to *r*_p_ = 3*σ*_PSF_ for all pinholes.

From Figures 5d and 5e, we see that the triangle configuration from Figure 5a results in a precision improvement in the *x*-direction of at most 1.94 compared to SMLM, when the distance between the pinholes and the center of the triangle is *r* = 2*σ*_PSF_. As seen for the two-pattern case, this optimum is a result of two contrasting factors. On one hand, increasing the pattern distance illuminates the emitter with the tail of the Gaussian intensity profile, thereby increasing the information that signal photons carry about the relative distance between the illumination pattern and the emitter. On the other hand, increasing the distance between the emitter and the pinholes also increases the amount of signal photons that are blocked by the spinning disk, while the pattern-dependent background increases due to the low illumination intensity.

**FIG. 5.**
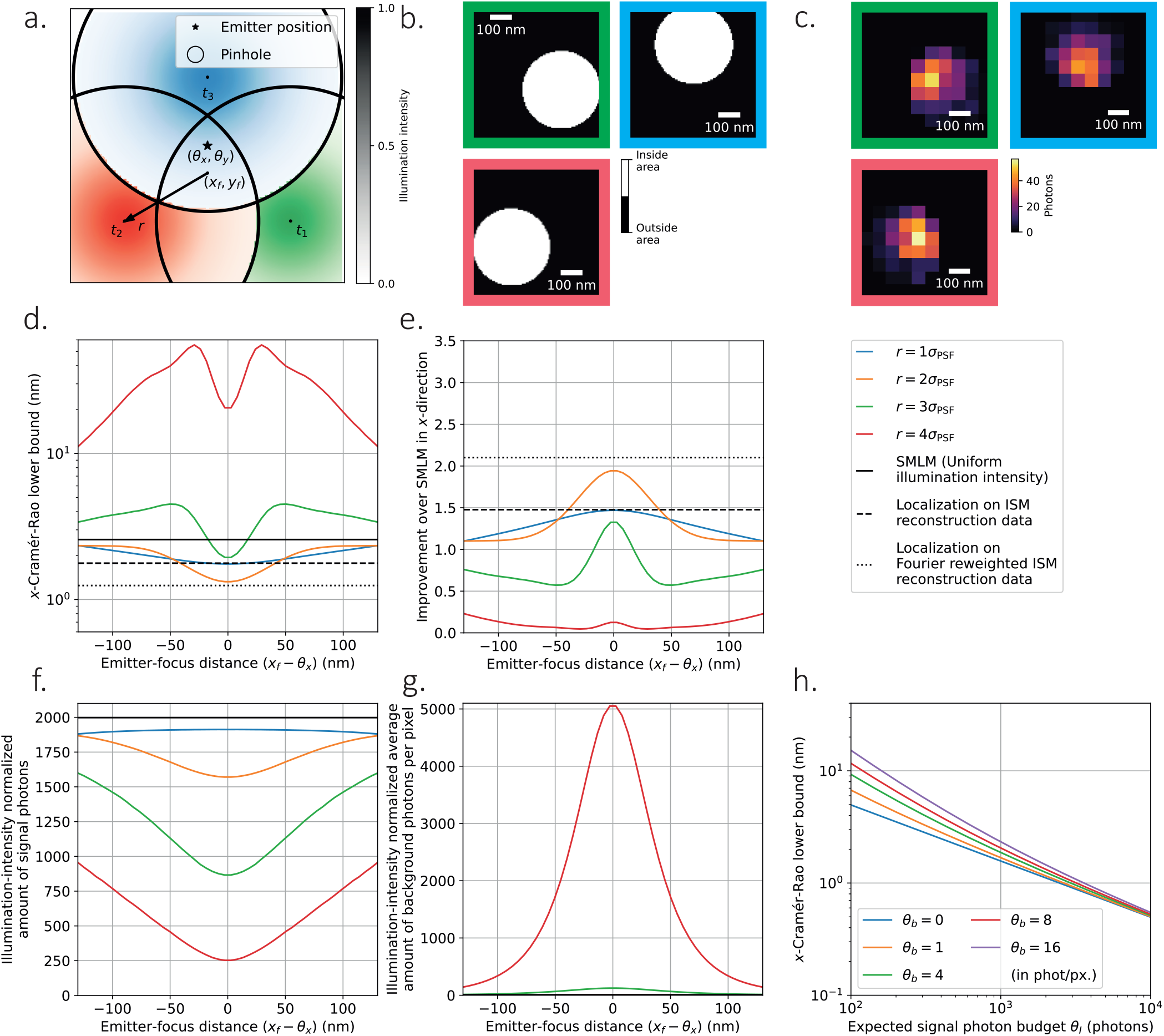
Theoretical minimum localization uncertainty of SpinFlux localization with three pinholes and patterns in an equilateral triangle configuration. In (c-g), we used 2000 expected signal photons and 8 expected background photons per pixel, with pinhole radius *r*_p_ = 3*σ*_PSF_. Results are evaluated for the scenario where the entire signal photon budget is exhausted after illumination with all patterns (disregarding signal photons blocked by the spinning disk). **(a)** Schematic overview of SpinFlux localization with a triangle of three pinholes, centered at focus coordinates (*x*_f_, *y*_f_). In (d-g), the *x*-distance (*x*_f −_*θ*_*x*_) between the pattern focus and the emitter is varied, where *y*_f_ = *θ*_*y*_. **(b)** Example of pinholes in the region of interest (650 ×650 nm). The pinhole radius *r*_p_ = 2*σ*_PSF_ and pinhole spacing *r* = 1.5*σ*_PSF_ were used. The pinhole masks were discretized with *N*_M,*x*_, *N*_M,*y*_ = 100 mesh pixels in each direction. **(c)** Example of fluorescent response in the region of interest, resulting from illumination and emission through each pinhole in (b). **(d)** Cramér-Rao lower bound (CRLB) in *x*-direction as a function of the emitter-focus *x*-distance. Simulations show SpinFlux with varying pinhole spacing and widefield single molecule localization microscopy (SMLM). **(e)** Improvement of the SpinFlux CRLB over SMLM as a function of the emitter-focus *x*-distance for varying pinhole spacing. **(f)** Average amount of signal photons after compensation for non-maximum illumination intensity as a function of the emitter-focus *x*-distance, for SpinFlux with varying pinhole spacing and widefield single molecule localization microscopy (SMLM). **(g)** Average amount of background photons per pixel after compensation for non-maximum illumination intensity as a function of the emitter-focus *x*-distance, for SpinFlux with varying pinhole spacing and widefield single molecule localization microscopy (SMLM). **(h)** CRLB in *x*-direction as a function of expected signal photon count for varying values of the expected background photon count. The pinhole radius *r*_p_ = 3*σ*_PSF_ and pinhole spacing *r* = 2*σ*_PSF_ were used and (*x*_f_, *y*_f_) = (*θ*_*x*_, *θ*_*y*_).

These effects are also present in the other scenarios under our consideration. In case the illumination power and time are kept constant, as in Supplementary Figure 16, the largest improvement in *x*-direction over SMLM is 1.09 for *r* = *σ*_PSF_, as the signal photon count decays sharply for larger spacing. When disregarding the effects of pattern-dependent background, shown in Supplementary Figure 26, the optimal spacing is *r* = 3*σ*_PSF_. This results in a precision improvement of 2.38 over SMLM.

Note that in all these cases, the *x*-localization precision of the triangle configuration is worse than that of the two-pattern configuration of Subsection III C. The reason for this is that the triangle configuration contains one pinhole, of which the *x*-coordinate is located close to the true emitter *x*-coordinate (i.e. the blue pattern in Figure 5a). As such, signal photons that are collected after illumination with this pattern contain little information about the emitter *x*-position. The two-pattern configuration of Subsection III C is thus able to distribute signal photons more efficiently, to maximize the information about the emitter *x*-position.

On the other hand, as discussed earlier for Supplementary Figures 6, 15 and 25, the two-pattern configuration contains little information about the emitter *y*-position. To investigate this for the triangle configuration, Supplementary Figures 7, 17 and 27 show the *x*-localization precision that can be achieved when the triangle pattern is rotated clockwise by 90 degrees for all three scenarios under consideration. Equivalently, these results also hold for the *y*-precision that can be attained with the non-rotated pattern. In all scenarios, it can be seen that the optimal spacing *r* and the localization precision are comparable to those for the non-rotated triangle configuration. Specifically for the scenario where the entire signal photon budget is exhausted, we find a precision improvement in *y*-direction of 2.05 over SMLM. As the rotated pattern is asymmetric along the *x*-axis, the precision also scales asymmetrically around the optimum. Additionally, the asymmetry causes a shift to the optimal *x*-coordinate of the pattern focus. For example, the optimal focus position is *x*_f_ = *θ*_*x*_ *−*0.13 nm when considering the scenario where the entire signal photon budget is exhausted.

From the equivalence, we find that the triangle configuration balances the localization precision in the *x*and *y*-directions at approximately a twofold improvement in either direction, at the cost of suboptimal precision in each individual direction.

In MINFLUX^8,15^, a triangle configuration was also used for illumination, where an additional fourth pattern was added in the center of the configuration. As such, we also consider the scenario where an additional pinhole and pattern are added in the center of the triangle for all three scenarios under consideration and for both rotations of the configuration (see Supplementary Figures 8, 9, 18, 19, 28, 29).

From Supplementary Figures 8 and 9, we find that adding a center pinhole causes a deterioration of the localization precision, compared to the triangle configuration without a center pinhole. Specifically for the scenario where the entire signal photon budget is exhausted, the precision improvement over SMLM is at most 1.44 for the non-rotated pattern, and at most 1.78 for the rotated pattern. On the other hand, the domain where SpinFlux attains an improvement over SMLM has increased due to the addition of the center pinhole. For the non-rotated pattern with spacing *r* = 2*σ*_PSF_, the improvement over SMLM varies between 1.39 and 1.44 as long as the pattern focus and the emitter remain in a 130 nm distance from each other.

The explanation for both these effects is that the center pinhole blocks the least amount of signal photons, and also claims the majority of the signal photon budget due to illumination with near-maximum intensity. As such, as shown in Supplementary Figures 8f, 8g, 9f and 9g, the effect of the pinhole spacing *r* on the usage of the signal photon budget and background count is strongly reduced. For pattern spacings between *r* = 0.5*σ*_PSF_ and *r* = 2*σ*_PSF_, pattern focus positions within a 130 nm range of the emitter position and either rotation, signal photon counts vary between 1753 and 1968 photons and average backgrounds vary between 0.88 and 4.30 photons per pixel. When the center of the triangle is displaced from the emitter position, another pinhole is able to cover the emitter position, thereby enlarging the range of similar photon counts and increasing the domain of precision improvement.

We find similar effects when neglecting the pattern-dependent background, as shown in Supplementary Figures 28 and 29. These can be explained by a similar reasoning for the signal photon count as above. However, we do not see this for the scenario where the illumination power and time are constant for each pattern (see Supplementary Figures 18 and 19). In this scenario, the patterns on the corners of the triangular configuration add little information due to their low illumination intensity. On the other hand, the center pattern illuminates with high intensity, resulting in a signal photon response that is comparable to the single-pattern case of Supplementary Figure 10. Overall in this scenario, the triangle configuration with a center pinhole is able to improve the precision over the same configuration without a center pinhole, with a maximum improvement of 1.14 over SMLM.

## IV. DOUGHNUT-SHAPED INTENSITY PATTERNS

Note that MINFLUX uses a doughnut-shaped intensity pattern for illumination, which contains an intensity minimum in the center. As described until now, SpinFlux uses a Gaussian intensity profile, with an intensity maximum in the center. By incorporating a phase mask in the illumination and emission paths (see Supplementary Figure 30), SpinFlux can be adapted to utilize doughnut-shaped illumination. As the doughnut-shaped pattern increases the information content of signal photons in its center rather than at its boundary^8^, it will mitigate the situation where highly informative signal photons are blocked by the pinhole, which in turn improves the theoretically minimum localization uncertainty. We explore this effect in Supplementary Figures 31 to 42.

Supplementary Figures 31 and 32 show the SpinFlux localization precision of the triangular configuration without a center pinhole, in the scenario where the entire signal photon budget is exhausted. Here, the improvement of SpinFlux with doughnut-shaped illumination over SMLM is approximately 1.64 in the *x*-direction and 1.74 in the *y*-direction at a pinhole spacing *r* = 3*σ*_PSF_. This improvement is comparable to that of SpinFlux with Gaussian illumination, as the intensity minimum of the illumination doughnut is placed 3*σ*_PSF_ away from the emitter. The Gaussian pattern at *r* = 2*σ*_PSF_ and the doughnut-shaped pattern at *r* = 3*σ*_PSF_ are comparable on the emitter coordinates, thereby negating the advantages of the doughnut-shaped pattern.

This changes when including a center pinhole in the triangular configuration, as shown in Supplementary Figures 33 and 34. Here, the maximum improvement over SMLM is 3.5 in the *x*and *y*-directions at a pinhole spacing of *r* = 4*σ*_PSF_. When increasing the spacing *r* between the pinholes (beyond the width of the doughnut-shaped beam), a larger share of the signal photon budget will be claimed by the center pinhole. The intensity minimum of the center pinhole increases the information content of signal photons, thereby improving the resolution over SpinFlux with Gaussian illumination. However, this improvement decays sharply when the pattern focus is not centered on the emitter position. Specifically for *r* = 4*σ*_PSF_, the improvement exceeds 1.5 in either direction only when the emitter-focus distance is smaller than 5 nm. Therefore, it is more practical to choose a smaller spacing between the pinholes. For *r* = 3*σ*_PSF_, the maximum improvement over SMLM is 3.3 in the *x*and *y*-directions, and the improvement is larger than 1.5 in either direction when the emitter-focus distance is at most 37 nm.

In the other scenarios under our consideration, we find similar conclusions. When considering the scenario where the illumination power and time are constant for each pattern (see Supplementary Figures 35 to 38), the triangular configuration with center pinhole improves the localization precision over SMLM by a factor 1.5 in the *x*-direction and by a factor 1.8 in the *y*-direction at a pinhole spacing of *r* = 0.5*σ*_PSF_. As the illumination energy is limited in this scenario, the information increase due to illumination through the center pinhole is balanced out by a lowered total signal photon response between 500 and 600 signal photons. When neglecting pattern-dependent background (see Supplementary Figures 39 to 42), the maximum improvement over SMLM is 19.6 in *x*and *y*-directions at a pinhole spacing of *r* = 4*σ*_PSF_.

## V. CONCLUSION

In meSMLM, sparse activation of single emitters with patterned illumination results in improved localization precision over SMLM. Additionally, meSMLM improves the resolution over image reconstruction in SIM while reducing the required amount of illumination patterns. As such, we need ways to enable meSMLM techniques in existing setups which are limited by image reconstruction in processing.

We developed SpinFlux, which incorporates meSMLM into SDCM setups. In SpinFlux, patterned illumination is generated using a spinning disk with pinholes to illuminate the sample with. Subsequently, the emission signal is windowed by the same pinhole before being imaged on the camera.

We have derived a statistical image formation model for SpinFlux, which includes the effects of patterned illumination, windowing of the emission signal by the pinhole and pattern-dependent background. For our analysis, we considered Gaussian illumination patterns and a Gaussian emission PSF. We also consider doughnut-shaped illumination patterns, which can be generated by incorporating a phase mask in the illumination path. In addition, we have derived and evaluated the CRLB for this model. We applied the CRLB to various illumination pattern configurations, to quantify the theoretical minimum uncertainty that can be gained with SpinFlux.

When using one pattern only, pattern-dependency of the background causes an improvement of at most 1.17 over SMLM, whereas no improvement is found when neglecting this effect. In the single-pattern case, the pinhole blocks signal photons which carry information about the emitter position. As such, it is beneficial for SpinFlux to use pinholes which are as large as possible to reduce the amount of signal photons blocked by the pinhole. In other words, we find that a spinning disk with pinholes is convenient to generate patterned illumination, although the pinhole itself has an adverse effect on the localization precision due to the blockage of signal photons.

However, we have not considered neighboring emitters in our analysis, nor have we modeled out-of-focus background. In SDCM, optical sectioning is achieved with the spinning disk by reducing the effects of neighboring or out-of-focus fluorescent signal, thereby improving the resolution. We expect that the pinhole has a similar effect on the localization precision that can be attained with SpinFlux, thereby resulting in an optimal pinhole radius. Future research should focus on incorporating these effects into the SpinFlux image formation model.

Based on the single-pattern results, we conclude that SpinFlux requires multiple patterns to generate a significant precision improvement over SMLM. We explored various multiple-pattern configurations. We found that a configuration of two pinholes with radius 3*σ*_PSF_, separated in *x*-direction around the emitter position by a distance of 4*σ*_PSF_, results in a precision improvement of 2.62 in *x*-direction compared to SMLM, while the *y*-improvement is at most 1.12. For larger separations, the information content of signal photons increases due to illumination with the low-intensity tails of the Gaussian illumination pattern. However, when the separation increases above 4*σ*_PSF_, the loss of signal photons due to the windowing effect of the pinhole causes deterioration of the localization precision.

We also evaluated the theoretical minimum uncertainty of a triangular pattern configuration, where pinholes are located at the corners of an equilateral triangle around the emitter position. This results in approximately a twofold *x*-precision improvement over SMLM, which is a reduction compared to the two-pattern configuration. However, the triangle configuration also attains approximately a twofold precision improvement in the *y*-direction. As such, the triangle configuration balances the localization precision in the *x*- and *y*-directions, at the cost of suboptimal precision in each individual direction. Including a center pinhole in the triangle does not improve the maximum localization improvement, but it stretches out the domain on which any improvement can be attained.

By including a phase mask in the illumination and emission paths, illumination patterns with arbitrary diffraction-limited intensity profiles can be created. We evaluated the localization precision of SpinFlux with doughnut-shaped illumination. As the doughnut-shaped pattern increases the information content of signal photons in its center rather than at its boundary, it will mitigate the situation where highly informative signal photons are blocked by the pinhole. We find that in the triangular configuration with a center pinhole, the maximum improvement over SMLM is increased to 3.5 in the *x*- and *y*-directions at a pinhole spacing *r* = 4*σ*_PSF_. We conclude that SpinFlux is an effective and versatile method to integrate meSMLM in existing SDCM setups. While localization on ISM data ideally results in an average global improvement of 1.48 over SMLM, or 2.10 with Fourier reweighting, SpinFlux is the method of choice for local refinements of the localization precision.

## SUPPORTING CITATIONS

References ^30,31^ appear in the Supplementary Material.

## SUPPLEMENTARY MATERIAL

See supplementary material for Supplementary Notes 1 to 4 on the derivation of the SpinFlux image formation model, the CRLB and the model partial derivatives needed to compute the CRLB for SpinFlux. Supplementary Figures 1 to 42 show additional schematics and results and Supplementary Table 1 lists simulation parameter values.

## DATA AVAILABILITY STATEMENT

The data that support the findings of this study are openly available in 4TU.ResearchData^32^ at https://doi.org/10.4121/21313230. The code that supports the findings of this study is openly available on GitHub^33^ at https://github.com/qnano/spinflux-crlb.

## ACKNOWLEDGMENTS

D.K., S.H. and C.S.S. were supported by the Netherlands Organisation for Scientific Research (NWO), under NWO START-UP project no. 740.018.015 and NWO Veni project no. 16761.

## AUTHOR’S CONTRIBUTIONS

D.K., S.H. and C.S.S. designed the research. D.K. derived and implemented the model, analyzed the data and wrote the manuscript, which was edited by S.H. and C.S.S. The study was supervised by C.S.S.

